# Dendro-plexing single input spikes by multiple synaptic contacts enriches the computational capabilities of cortical neurons and reduces axonal wiring

**DOI:** 10.1101/2022.01.28.478132

**Authors:** David Beniaguev, Sapir Shapira, Idan Segev, Michael London

## Abstract

A cortical neuron typically makes multiple synaptic contacts on the dendrites of its postsynaptic target neuron. The functional implications of this apparent redundancy are unclear. Due to dendritic cable filtering, proximal dendritic synapses generate brief somatic postsynaptic potentials (PSPs) whereas distal synapses give rise to broader PSPs. Consequently, with multiple synaptic contacts, a single presynaptic spike results in a somatic PSP composed of multiple temporal profiles. We developed a “Filter-and-Fire” (F&F) neuron model that incorporates multiple contacts and cable filtering; it demonstrates threefold increase in memory capacity as compared to a leaky Integrate-and-Fire (I&F) neuron, when trained to emit precisely timed spikes for specific input patterns. Furthermore, the F&F neuron can learn to recognize spatio-temporal input patterns, e.g., MNIST digits, where the I&F model completely fails. We conclude that “dendro-plexing” single input spikes by multiple synaptic contacts enriches the computational capabilities of cortical neurons and can dramatically reduce axonal wiring.

## Introduction

In recent decades it has been shown experimentally that when cortical neurons are synaptically connected they typically connect to each other via multiple synaptic contacts rather than a single contact^1–5^. Multiple synaptic contacts originating from a single presynaptic axon often impinge on different parts of the dendritic tree of the post-synaptic neuron both in rodents ^1–3^ and also in humans^6,7^. Interestingly, if the formation of synaptic contacts were based purely on the proximity of the axon-to-the-dendrite and independent of each other (“Peters’ rule”^8^) then one would expect that the distribution of the number of such multiple contacts would be geometric, with one contact per axon being the most frequent case ^9–11^. This is far from what was empirically observed where, e.g., around 4-8 synaptic contacts are formed between pre- and postsynaptic layer 5 cortical pyramidal neurons^5^. This deviation from the distribution predicted by Peters’ rule suggests that the number of synaptic contacts between two connected neurons is tightly controlled by some developmental process and is thus likely to serve a functional purpose.

Several phenomenological models have attempted to explain how multiple synaptic contacts between the presynaptic and postsynaptic neurons are formed^9^, but only a few studies have attempted to tackle the question of how they might be beneficial from a computational perspective. It is typically thought that this redundancy overcomes the problem of probabilistic synaptic vesicle release, which results in unreliable signal transmission between the connected neurons^12^. However, the same effect using a simpler mechanism could be achieved by multiple vesicles release (MVR) per synaptic activation^1,12^ and does not require multiple synaptic contacts (which is more “expensive” to establish). Hiratani and Fukai^13^ demonstrated that multiple synaptic contacts might allow synapses to learn quicker. However, faster learning, although beneficial, fundamentally does not endow the neuron with the ability to perform new kinds of tasks. Recently, Zhang et al.^14^ incorporated multiple contacts in the context of deep artificial neural networks but did not demonstrate tangible computational benefits. Jones et al.^15^ also model dendrites in the context of artificial neural networks demonstrating that, when using a threshold linear dendrite model, the classification performance of a single neuron with multiple contacts is improved compared to the single contact case. Several other studies use multiple synaptic contacts in the context of artificial neural networks, demonstrating some computational benefits^16–18^.

In the present study we propose two key functional consequences of multiple contacts. Towards this end, we developed a simplified spiking neuron model that we termed the Filter & Fire (F&F) neuron model. This model is based on the Integrate and Fire (I&F) neuron model^19,20^ but incorporates two key additional features: (i) It takes into account the effect of the dendritic cable filtering on the time course of the somatic PSP, whereby proximal synapses generate brief somatic PSP while identical synapses that are located at distal locations on the dendritic tree give rise to broad somatic PSPs^21,22^; (ii) Each presynaptic axon makes multiple synaptic contacts on the F&F neuron model. Consequently, a single presynaptic spike results with PSP that is composed of multiple temporal profiles (we term this phenomenon “dendro-plexing” of the presynaptic spike).

To analyze the memory capacity of this model, we used the formulation of Memmesheimer et al.^23^ developed for the I&F model. In this approach, the model is trained (via changes in synaptic strengths) to emit precisely-timed output spikes for a specific random input pattern; the capacity is defined as the maximal number of precisely timed output spikes during some time period divided by the number of incoming input axons. We show that the memory capacity of the F&F model is three-fold larger than that of the I&F model. We next show the F&F neuron can solve real-world classification tasks where the I&F model completely fails. We further explored the effect of unreliable synapses on the memory capacity of the F&F model and, finally, we demonstrate that multiple synaptic contacts dramatically reduce axonal wiring requirement in cortical circuits.

## Results

### Mathematical description of the filter and fire (F&F) neuron model with multiple synaptic contacts

We propose hereby a Filter and Fire (F&F) neuron model, which is similar to the standard current-based Leaky Integrate and Fire (I&F) spiking neuron model, but with two added features. The first feature incorporates the temporal characteristics of a dendritic cable as initially demonstrated by Rall (Rall 1964; Rall 1967) in which, due to cable filtering, synaptic inputs that connect at distal dendritic locations exhibit prolonged PSPs at the soma (Fig. 1**B** top traces), whereas proximal synaptic inputs generate brief PSP profiles (Fig. 1**B** bottom traces). The second feature is that each input axon connects to multiple locations on the dendritic tree, sometimes proximal and sometimes distal (Fig. 1**A,B**).

**Fig. 1.**
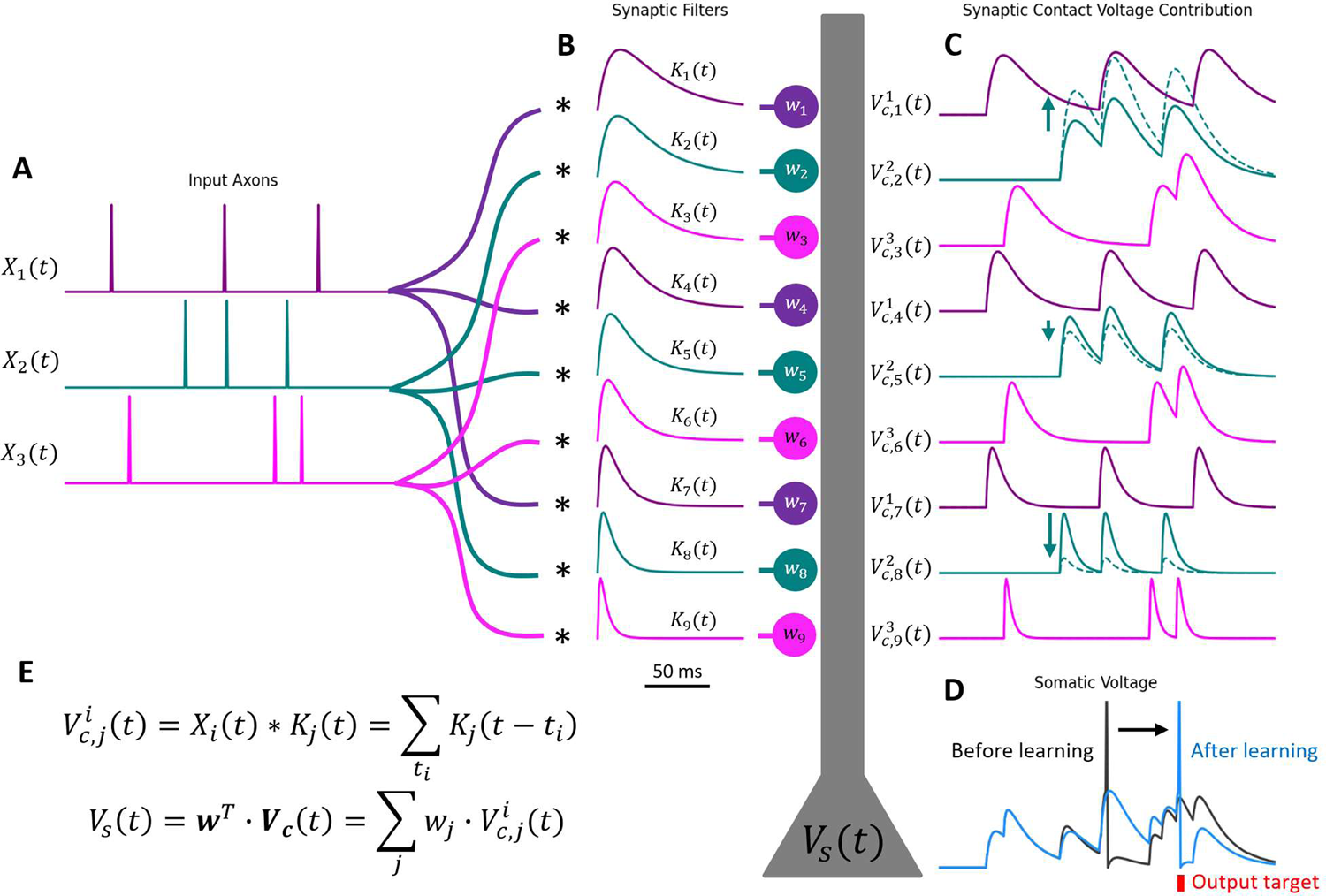
The Filter and Fire (F&F) neuron receives input through multiple synaptic contacts per axon and filters each contact with a different synaptic kernel (“dendritic multiplexing”) (A) Example for three incoming input axons, each making three synaptic contacts onto the postsynaptic neuron. (B) Various synaptic filters (kernels) representing their respective locations on the dendritic tree (shown schematically in grey). Proximal synaptic filters are brief, whereas more distal synaptic filters have broader temporal profiles. Kernel colors are according to the source axon. (C) Individual contact voltage responses at the synaptic loci that result from the convolution (* symbol) of an axonal spike train with the respective synaptic filter. Continuous line is the voltage contribution of the contact before learning and dashed lines after learning. In this example, the weight of the second-from-top green synaptic contact, *W*_z_, was increased, whereas *W*_S_ and *W*_8_ decreased following learning. The weight of each contact can change independently during learning. (D) Somatic voltage is a weighted sum of the contributions of each synaptic contact. In black is the somatic trace before learning and in blue after learning. Standard I&F reset mechanism was applied for the spike generation mechanism at the soma. (E) Equations formally describing the synaptic contact voltage contributions and the somatic voltage.

Formally, consider *N*_axons_ the number of input axons (Fig. 1**A**), denoted by index i, and their spike trains denoted by *X*_i_(*t*). Each axon connects to the dendrite via *M* synaptic contacts (*M* = 3 is illustrated in Fig. 1). Each contact connects to the dendrite at a location denoted by index j and filters the incoming axon spike train with a specific synaptic kernel *K*_j_(*t*). The voltage contribution trace (PSP) of this contact is: *V*^i^ (*t*) = *X*_i_(*t*) ∗ *K*_j_(*t*) = ∑_t_ *K*_j_(*t* − *t*_i_). There is a total of *M* · *N*_axons_ such contacts and therefore the same number of overall synaptic contact voltage contribution traces (Fig. 1**C**). In vector notation we denote *V*_c_(*t*) = [*V*^1^ (*t*), *V*^z^ (*t*), ⋯, *V*^Naxons^ (*t*)]. Each synaptic contact has a weight, *W*. In vector notation, *W* = [*W*_1_, *W*_z_, ⋯, *W*_M·Naxons_]. The contribution of each contact is multiplied by its corresponding weight to form the somatic voltage trace: *V*_s_(*t*) = *W*^T^ · *V*_c_(*t*) = ∑_j_ *W*_j_ · *V*_c,j_(*t*) (Fig. 1**D**). When the spike threshold is reached, a standard reset mechanism is applied. Please note that the “dendrites” in this model are linear and therefore retain the analytic tractability of the I&F neuron model.

Here we model the temporal ramifications of the effect of adding a passive dendritic cable; we did not consider the effect of nonlinear dendrites (see Discussion). The kernels we used are typical double exponential PSP shapes of the form: *K*_j_(*t*) = *A* · (*e*^-t⁄rdecay,j^ − *e*^-t⁄rrise,j^), where A is a normalization constant such that each filter has a maximum value of 1, and τ_decay,j_, τ_rise,j_ are randomly sampled for each synaptic contact, representing randomly connected axon-dendrite locations. Note that for mathematical simplicity we do not impose any restrictions on synaptic contact weights, each weight can be both positive or negative regardless of which axon it originates from. Indeed, the goal of the study is not to replicate all particular biological details, but rather to specifically explore the computational benefit that arises due to two key general dendritic/connectivity features: temporal filtering of synaptic potentials due to dendritic cable properties and multiple synaptic connections between pairs of cortical neurons.

### Increased memory capacity of the F&F neuron with multiple synaptic contacts

We first tested the memorization capacity of the F&F neuron model as a function of the number of multiple connections per axon. We utilized the framework proposed by Memmesheimer et al.^23^ to measure memory capacity, and used their proposed local perceptron learning rule for that task. In short, this capacity measure indicates the maximal number of precisely timed output spikes in response to random input stimulation during some time period divided by the number of input axons. Fig. 2**A** shows random spiking activity of 100 input axons for a period of 60 seconds (top). Below the output of the postsynaptic neuron is shown before learning (black), after learning (blue) and the desired target output spikes (red). In this example, we used *M* = 5 multiple contacts per axon. For the given set of random input spike trains, it is possible to find a synaptic weight vector to perfectly place all output spikes at their precisely desired timing. In Fig. 2**B** we repeat the simulation shown in Fig. 2**A** for various values of multiple contacts (M) while re-randomizing all input spike trains, desired output spike trains, and the synaptic filter parameters of each contact. We repeat this both for the I&F neuron model (i.e., a single synaptic PSP kernel for all synapses/axons) and the F&F neuron model with a randomly selected synaptic kernel for each synapse (see **Methods** for full details of the kernel shapes used). The y axis represents our success in placing all of the output spikes accurately, as measured by the area under the receiver operating characteristic (ROC) curve (AUC) for the binary classification task of placing each spike in 1ms time bins. Error bars represent the standard deviation of the AUC over multiple repeats (18), while re-randomizing the input, re-randomizing the synaptic kernels and re-randomizing the desired output spike trains.

**Fig. 2.**
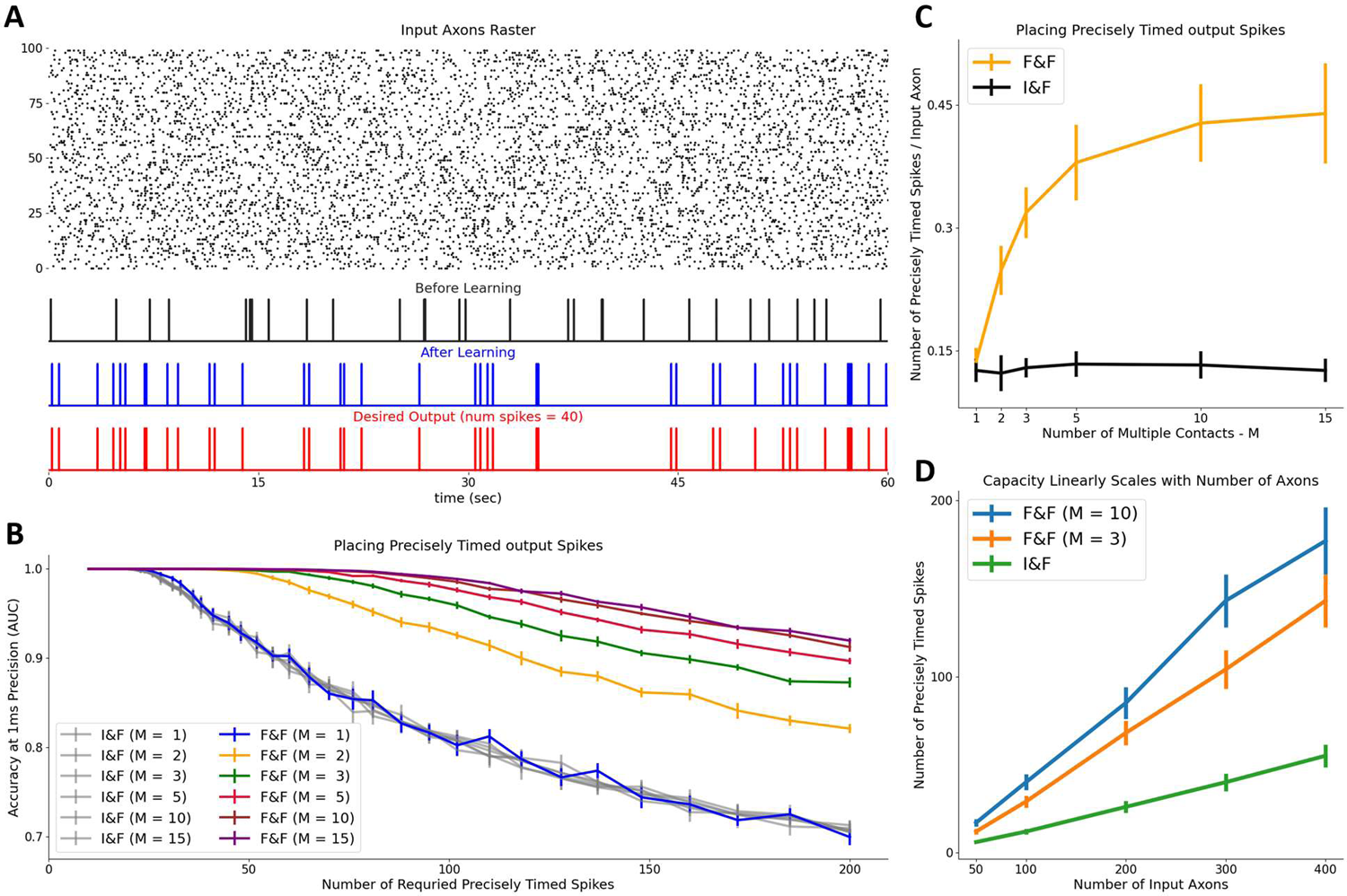
Increased memory capacity of precisely timed output spikes of the F&F neuron versus the I&F neuron. (**A**) Learning to place precisely timed output spikes for randomly generated input with F&F neuron model with M=5 multiple contacts. **Top**. Random axonal input raster plot. **Bottom**. Output spikes of the F&F model before learning (top, black), after learning (middle, blue) and the desired output spikes (bottom, red). (**B**) Binary classification accuracy at 1ms temporal resolution as measured by area under ROC curve (AUC) as a function of the number of required output spikes for input with 200 input axons. The capacity of the F&F models increases as the number of multiple contacts increases, whereas no such increase is observed for the I&F case. (**C**) Capacity as a function of the number of multiple connections per axon. For this plot we use the maximal number of spikes that achieves accuracy above AUC threshold of 0.99. The vertical axis depicts the fraction of successfully timed spikes for each input axon. Note that the saturation in the capacity for many multiple contacts is ∼3x compared to the respective I&F capacity. (**D**) The capacity scales linearly as a function of the number of input axons and exhibits no saturation.

Fig. 2**B** shows that all the curves obtained from the I&F neuron models cluster together, independent of M (the number of multiple contacts). This is to be expected as this is the classic case of synaptic redundancy when using a single temporal kernel for all presynaptic axons. For example, for a single axon and two contacts one can see that, for the case of a single synaptic kernel (as is the case in I&F model), the somatic voltage can be written as *V*_s_(*t*) = *W*_1_ · ∑ *K*(*t* − *t*_i_) + *W*_z_ · ∑ *K*(*t* − *t*_i_) = (*W*| |1 + *W*| |2) · ∑ *K*(*t* − *t*_i_) = *W*_eff_ · ∑ *K*(*t* − *t*_i_). Meaning, that the different weights *W*_1_, *W*_z_ associated with the same input axon are redundant and equivalent to a single effective weight *W*_eff_, and therefore this difference is not utilized in the case of an I&F neuron model where all synaptic kernels are identical.

The case of the F&F neuron with *M* = 1, is identical to the case of an I&F model only that it has different kernels for different synapses. This change on its own does not affect the capacity of the model, as the number of learnable and utilizable parameters is identical to the I&F case with *M* = 1, and thus this curve lies with the other curves of the I&F models. However, for the F&F models with multiple contacts (*M* = 2,3,5,10,15) there is an increase in accuracy, demonstrating that some of the additional synaptic weights are utilized (Fig. 2B). Fig. 2**C** displays the maximal number of output spikes that can be precisely timed as a function of M, for both I&F and F&F models. We measure the number of precisely timed spikes as the maximal number of spikes that is above a high accuracy threshold (AUC > 0.99), normalized by the number of axons. This provides the number of precisely timed output spikes per input axon on the y-axis. The figure shows that, for the I&F model, the capacity is around 0.15, whereas for the F&F neuron with a large number of synapses per axon, it saturates at about 0.45 (∼3-fold increase in capacity). Already for 3-5 contacts/axon (as is the case found for connected pairs of cortical pyramidal neurons) the capacity approaches the saturation level (see **Discussion**). Note that the number of degrees of freedom (tunable parameters) scales linearly with the number of multiple contacts, so there is no obvious explanation for the observed saturation in the F&F model. We will explain the origin of this 3-fold increase compared to the I&F model in Figure 4.

In Fig. 2D we vary the number of input axons and observe linear scaling of the number of precise spikes achieved with different slopes for different number of multiple connections. This indicates that increasing the number of multiple connections increases the effective number of parameters utilized per axon. To better illustrate the results of Fig. 2 more intuitively, we show in Fig. S1**A** a simple case whereby the I&F neuron can emit temporally precise output spikes by employing a spatial strategy, effectively selecting the input axon traces that happen to correlate with the desired output trace. In Fig. S1**B** we show how a F&F neuron can employ a temporal strategy by weighting differently the synaptic kernels with different time scales, allowing it to select individual input spikes that correlated with the desired output spikes instead of the entire axonal spiking activity.

### The F&F neuron can learn real-life spatio-temporal tasks that the I&F neuron cannot

Next, we demonstrate new capabilities of the F&F neuron model with multiple synaptic connections that are beyond that of the I&F neuron model. For this purpose, we developed a new spatio-temporal task derived from MNIST task, which is a large database of handwritten digits ^24^. Towards this end, we converted the horizontal spatial dimension of the image (image width) into a temporal dimension (Fig. 3**A** top) with a uniform time warping, such that 20 horizontal pixels will be mapped onto T milliseconds. T is the duration of presentation of the respective pattern (say digit “3”). The vertical spatial dimension of the image (image height) is simply replicated five times so that 20 vertical pixels will be mapped onto 100 axons. We then sample spikes for each axon according to the time varying Poisson instantaneous firing rate with additional background noise. For example, in this way, an axon corresponds to a row in a handwritten digit image, and this axon fires spikes with increased probability at times when the pixels in that row are active. An example of the resulting input spike trains representing different digits is shown by the raster plot in Fig. 3**A** middle frame.

**Fig. 3.**
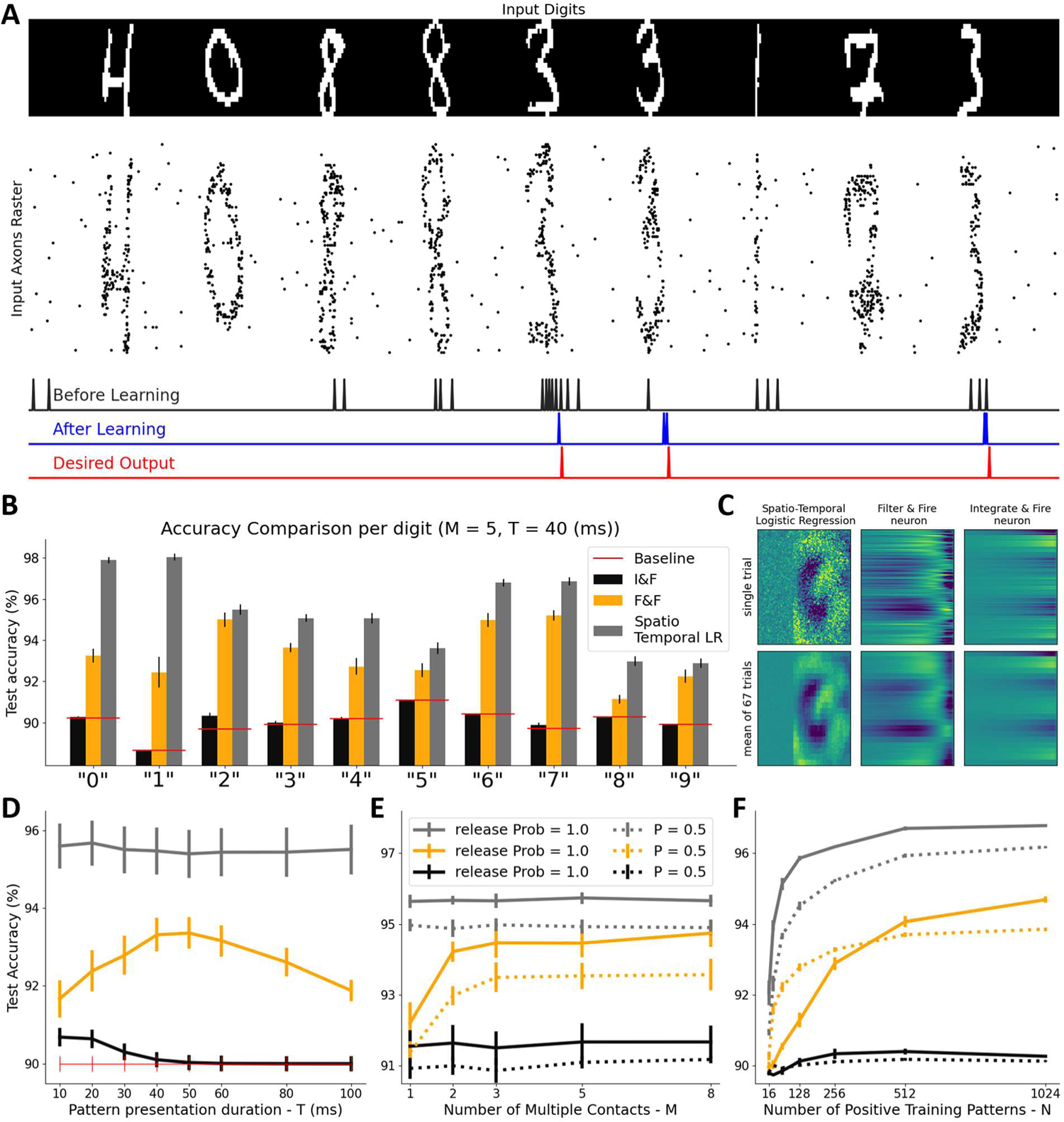
The F&F neuron can learn to recognize spatio-temporal patterns whereas the I&F neuron cannot. (**A**) A F&F neuron was trained to recognize digits in a spatio-temporal version of the MNIST task. Top: Original input digits; Middle: Raster plot of the axonal spikes (100 axons, 5 contacts per axon with duration of digit representation, T = 40 msec) representing the respective digits (with some additional background noise, see **Methods**). Bottom: Output of the F&F neuron (with 5 multiple contacts) before learning (black), after learning, (blue) and the desired output spike train (red). In this case the neuron was trained to detect the digit “3”. The depicted traces are from a new test set. (**B**) Full test set classification accuracy of the F&F neuron (orange), the I&F neuron (black) and a non-biologically plausible temporally sliding spatio-temporal Logistic Regression model (gray) for the spatio-temporal digit classification task, when each digit was used as the positive class and the rest of the digits were considered the negative class. The random chance baseline for each case is shown by the red horizontal lines (baseline fluctuates near 90% due to slightly unequal number of images in the MNIST dataset for each digit). The I&F models cannot learn the task whereas the F&F neurons sometimes approach the (non-biological) spatio-temporal logistic regression model. (**C**) Spatio-temporal representation of the learned weights of the three models when attempting to visually detect the digit “3”. The digit “3” can be clearly discerned in the logistic regression case, a faint “3” in the F&F case, whereas for the I&F model, an attempt to use spatial-only information to detect the digit is visible and the detection of the digit is poor, at chance level. The top three images are for a single trial with a single random axonal wiring, and Poisson sampling of the inputs. Bottom three images are an average of 67 trials. (**D**) Summary of test accuracy, averaged across all digits, for the three models as a function of the temporal duration (presentation time) of the digit. Time between digit presentations was 70 ms. The decay time constant of the synaptic kernel for the I&F model (black curve) was 30 ms, as is the maximal decay time constant for all synaptic kernels for the F&F model. The performance of the F&F neuron peaks at T = 40-50 ms (orange curve). In this case, the F&F model shown had M=5 multiple contacts. (**E**) Test accuracy as a function of the number of multiple contacts for all 3 models. The dashed line is the accuracy under the unreliable synapse’s regime with a release probability of 50% per contact. Note the consistent drop in accuracy. In this graph the positive digit used was “7” and the pattern temporal duration was T=30 ms. The number of positive training samples used in this case was subsampled to 2048 patterns. (**F**) Test accuracy as a function of the number of training samples. Each training sample was shown to the model 15 times and for each spike and each contact an independent probability of release was applied. Note the regularization benefit of unreliable synaptic release for a small number of training samples (dashed orange line above solid orange line).

Next, we trained the F&F neuron to produce a spike at the end of a specific digit. We trained the model on the full MNIST train subset of digits, and present results on an unseen test set. Before learning (black), after learning (blue), and the desired output (red) are presented at the bottom of Fig. 3**A** for the case where the selected digit to be recognized was “3”. We then repeat this process for all digits for three models - I&F neuron, F&F neuron, and a spatio-temporal, temporally sliding, logistic regression (LR) model ^25^. We note that the LR model is not biologically plausible and cannot be considered as a model of a neuron. Importantly, the LR model has *N*_axons_ · *T* learnable and fully utilizable parameters (*N*_axons_ parameters for each 1ms time bin) which is much greater than *N*_axons_ · *M* parameters that are used in the F&F and I&F models (which are also not fully utilizable as we have seen in Fig. 2).

The MNIST training set consists of 60,000 images, approximately 6,000 for each digit. The positive class for each digit classification in the experiments in Fig 3 consisted of 6.000 images, and the negative class around 54,000 images. When attempting to detect a single digit the baseline accuracy of a neuron that never spikes is around 90% accuracy, we therefore measure our accuracies compared to that baseline. The test accuracies following training for all models are depicted in Fig. 3**B**. For this plot a successful true positive (hit) is achieved if at least 1 spike has occurred in the time window of 10 ms around the ground truth desired spike. The temporal duration of each pattern T was 40 ms and the number of multiple contacts M was 5, for the plot in Fig. 3**B**. This figure clearly shows that the I&F neuron model is at chance level for almost all digits and is incapable of learning the task. In contrast, the F&F model is consistently better than chance, and sometimes approaches the “aspirational” spatio-temporal logistic regression model. In Fig. 3**C** we visually display the learned weights of all models when attempting to learn the digit 3. The weight matrix of the logistic regression model clearly depicts what appears to be an average-looking digit 3. The F&F neuron model depicts a temporally smoothed version of the logistic regression model, and the I&F model clearly cannot learn temporal patterns and therefore cannot recognize this digit at above chance level. For precise details of how the weights for the F&F and I&F models were visualized, see **Methods**. For a more simplified pattern classification case, see Fig. S1**C**. Fig. S1**D** shows how a F&F neuron can solve the task shown in Fig. S1**C**. Fig. S1**E** explains why this task cannot be solved by an I&F neuron.

The effect of T, the presentation duration of each digit, can be seen in Fig. 3**D**, which displays summary statistics of test accuracy, averaged across all digits, for the three models as a function of T. The interval between presented patterns was 70 ms and the decay time constant for the I&F model was 30 ms, to match the maximal decay time constant for all synaptic kernels in the F&F model. Fig. 3**D** shows that there is an optimal pattern presentation duration for the F&F model in the range of 40-50 ms, which is ∼1.5 times the maximal decay time constant in the model.

Next, we explored the effect of unreliable synaptic transmission and its interaction with multiple synaptic contacts. As explained in the **Introduction**, multiple synaptic contacts are often considered as a mechanism to overcome synaptic transmission unreliability. In Fig. 3**E** we display the test accuracy as a function of the number of multiple contacts for all 3 models for fully reliable synapses that we have displayed so far; the dashed line shows the accuracy under the unreliable synapse regime with release probability of p = 0.5 per contact. Note the consistent drop in test accuracy that does not go away even with a large number of multiple contacts. In this graph, the positive digit used was “7” and T = 30 ms. The number of positive training samples used in this case was 2048 patterns.

In principle, unreliable synaptic transmission could be considered as a mechanism for implementing “drop-connect”, a method used in training artificial neural networks that has a known regularization effect ^26,27^. In Fig. 3**F** we test if this is also applicable in our case. We show the test accuracy as a function of the number of training samples and demonstrate a regularization effect in the case of F&F that increases test accuracy for a low number of training input patterns. Note that each training sample was shown to the model multiple times (15 times in this case), and for each spike and each contact an independent probability of release was applied, effectively resulting in 15 noisy patterns that were presented to the neuron during training for each original training pattern. This suggests that unreliable synapses can also be viewed as a “feature” rather than a “bug” for the regime of a small number of training data points and can help avoid overfitting (see **Discussion**).

### Maximal capacity is explained by the effective 3-dimensional subspace spanning all synaptic kernels

Next, we wish to pinpoint the mathematical origin of the properties depicted in Figures 2 and 3 of the proposed F&F neuron model with multiple synaptic connections. We observed that all the PSP kernels we used have similar shapes and therefore the PSPs generated by the same axon will produce correlated inputs at the local synaptic responses vector *V*_c_(*t*). Any input correlation will limit the number of degrees of freedom available for learning by modification of the synaptic weights *W*. Fig. 4**A** shows all the PSPs used as heatmaps organized according to increasing values of τ_rise_ within each block and increasing τ_decay_ between the blocks. In Fig. 4**C** we show all the PSPs as temporal traces overlayed on each other. Both Fig. 4**A** and 4**C** clearly show that the shapes of the PSP kernels, although selected randomly and have some variance, are overall very similar to each other. We therefore apply singular matrix decomposition (SVD) on all PSP shapes (Fig. 4**B**) and found that 99.93% of the variance in all PSP shapes is explained by the first three singular vectors (Fig. 4**E**). This means that all synaptic kernels depicted in Fig. 4A and 4C are effectively spanned by a basis set of three orthogonal PSP-like shapes. For the sake of presentation and to avoid negative values in the trace shapes, we display the three independent kernels that are the result of non-negative matrix factorization (NMF) in Fig. 4**D**.

It is easy to see that these PSP shapes basically filter the input signal with various time constants and various delays. These are very intuitive shapes that we can easily interpret. This enables one to understand both the temporal smoothing aspect of the learned weights by the F&F neuron model (Fig. 3**C**) and also suggesting that the number of independent PSP shapes that span al the PSPs is a good candidate to explain the 3-fold increase in capacity shown in Fig. 2**C**. To verify this, we repeat the same experiment as in Fig. 2, but now each axon is connected to the post synaptic neuron via only three multiple connections, but this time we use the optimal PSPs shapes depicted in Fig. 4**D**. These results are depicted in Fig. 4**F**. The model with three orthogonal kernels has effectively identical results to those of the F&F neuron with randomly selected filters. First, to explain the resultant capacity, when we randomly sample more and more connections from the set of all possible kernels (e.g., increasing M), we slowly approach to span the entire three-dimensional space by each contact and therefore we have a saturation effect at three times the I&F baseline level. This means that we have reached the three independent kernels, each corresponding to an independent learnable parameter. Second, the specific shapes of these kernels explain the temporal smoothing effect we have observed in the learned weights of the F&F neuron model in Fig. 3**C**. Importantly, although we used somewhat artificial double exponential synaptic kernels, we verified that our results also hold for PSPs in a simulation of a highly realistic detailed L5 cortical neuron model. Fig. S2, shows that, despite some quantitative differences, qualitatively the results presented in our work using a reduced F&F neuron model are also valid for the case of a neuron with a complex dendritic tree receiving synaptic inputs over its entire dendritic tree. Namely, a basis set of three temporally distinct kernels can effectively span all synaptic kernels also in realistic models of cortical pyramidal neurons.

**Fig. 4.**
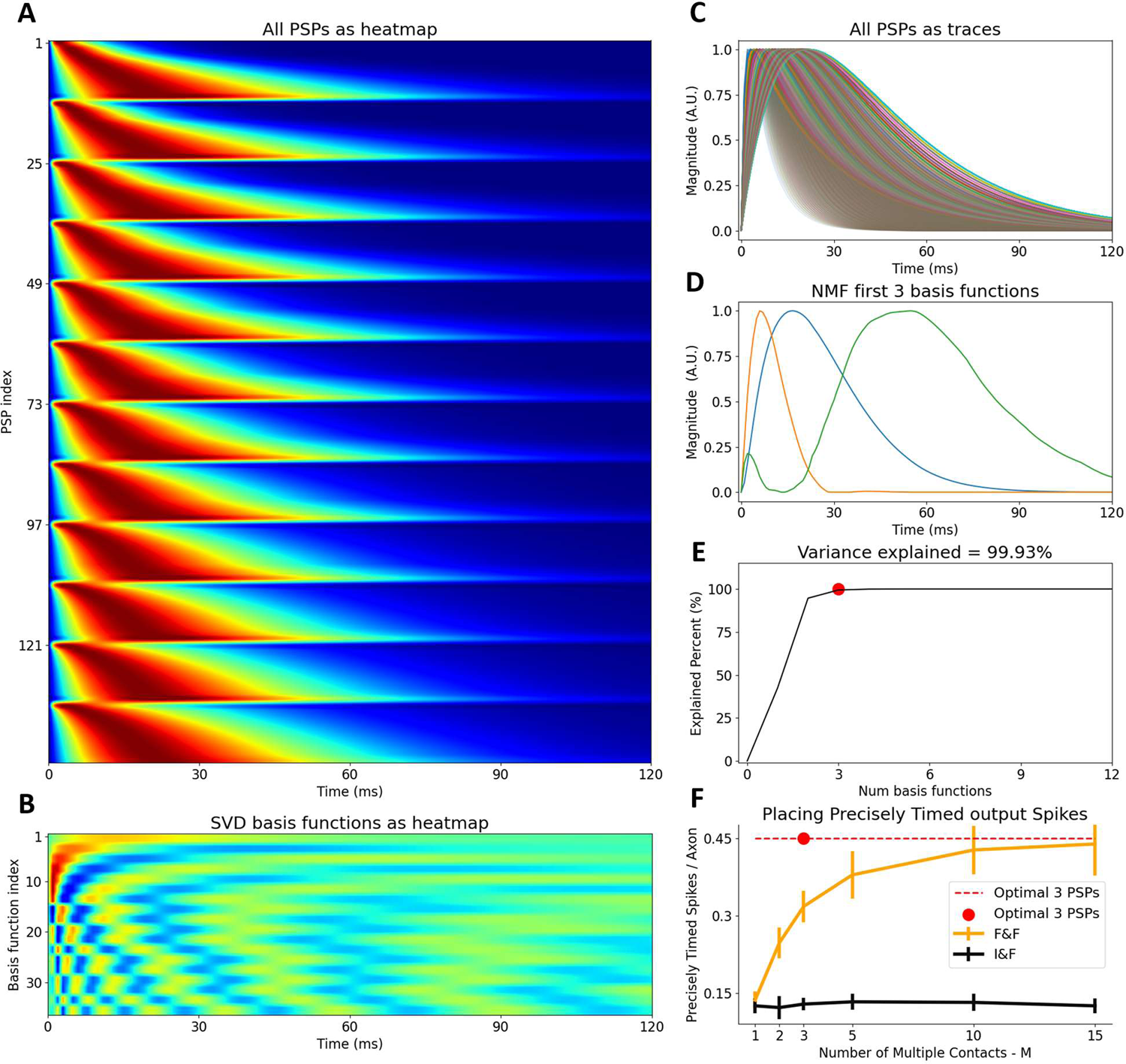
The dendritic filters are spanned by a three-dimensional basis set of PSPs accounting for the three-fold increase in the F&F capacity for a large number of multiple contacts as compared to the I&F model. (A) All possible postsynaptic potentials (PSPs) that were used in this study shown as heatmaps. Every row has a different rise time and decay time. (B) The Singular Value Decomposition of all the PSPs shown in (A) as heatmaps. This provides a Fourier-like basis set. (C) Traces of all post synaptic potentials (PSPs) that were used in the study. (D) First three basis functions of the non-negative matrix factorization (NMF) of all PSPs shown in (A & C). NMF instead of SVD was used for ease of interpretation and visualization purposes (see Methods). (E) Cumulative variance explained by each basis component. The three basis functions shown in (D) can span all PSPs shown in (C), red circle. (F) Direct verification that the three orthogonal basis set in (D) can be used as three optimal multiple contact filters achieving the same capacity as do 15 randomly selected multiple contacts.

### Multiple dendritic synapses per axon significantly reduce axonal wiring or network size

Fig. 5 shows that multiple dendritic synaptic contacts per axon allow cortical circuits to transmit axonal information in a more hardware-efficient way, by relying on downstream dendritic decoding. Three computationally equivalent alternatives are depicted in Fig 5**C,D**&**E**. Fig. 5**C** illustrates the case in which N axons transmit information that is readout by a downstream F&F neuron model with multiple synaptic contacts. In Fig, 5**D**, we illustrate a scenario of an equivalent I&F neuron. If we wish for a downstream I&F point neuron to produce the same output as that of the F&F in Fig. 5**C**, the spikes present in the N axons the F&F receives should spread out to 3N axons for the I&F neuron to receive as input (the “spreading” could even be random, as long as the information content is kept identical). This will allow the I&F in Fig.5**D** to precisely replicate the I/O transformation of the F&F in Fig.5**C**. Alternatively, each of the N axons could be replicated three times, each with some additional time delay, to account for the temporal dendritic integration properties implemented by the F&F neuron model and then fed to an I&F neuron. This last scenario is illustrated graphically in Fig. 5**E**. It is also verified in simulations in Fig. 5**A** for learning to produce precise spike time output and in Fig. 5**B** for solving the spatio-temporal MNIST digit recognition task. The three equivalent alternatives shown in Fig. 5**C, D****&E**, highlight the general statement that a system can tradeoff between putting recourses into encoding or decoding. In this particular case the three alternatives are (a) an encoder with 3N transmitting axons + 1 simple I&F neuron decoder (b) simple encoder with N transmitting axons + 3N simple delay decoding neurons + 1 simple I&F final decoder neuron and (c) simple encoder with N transmitting axons + 1 complex F&F decoder neuron.

**Fig. 5.**
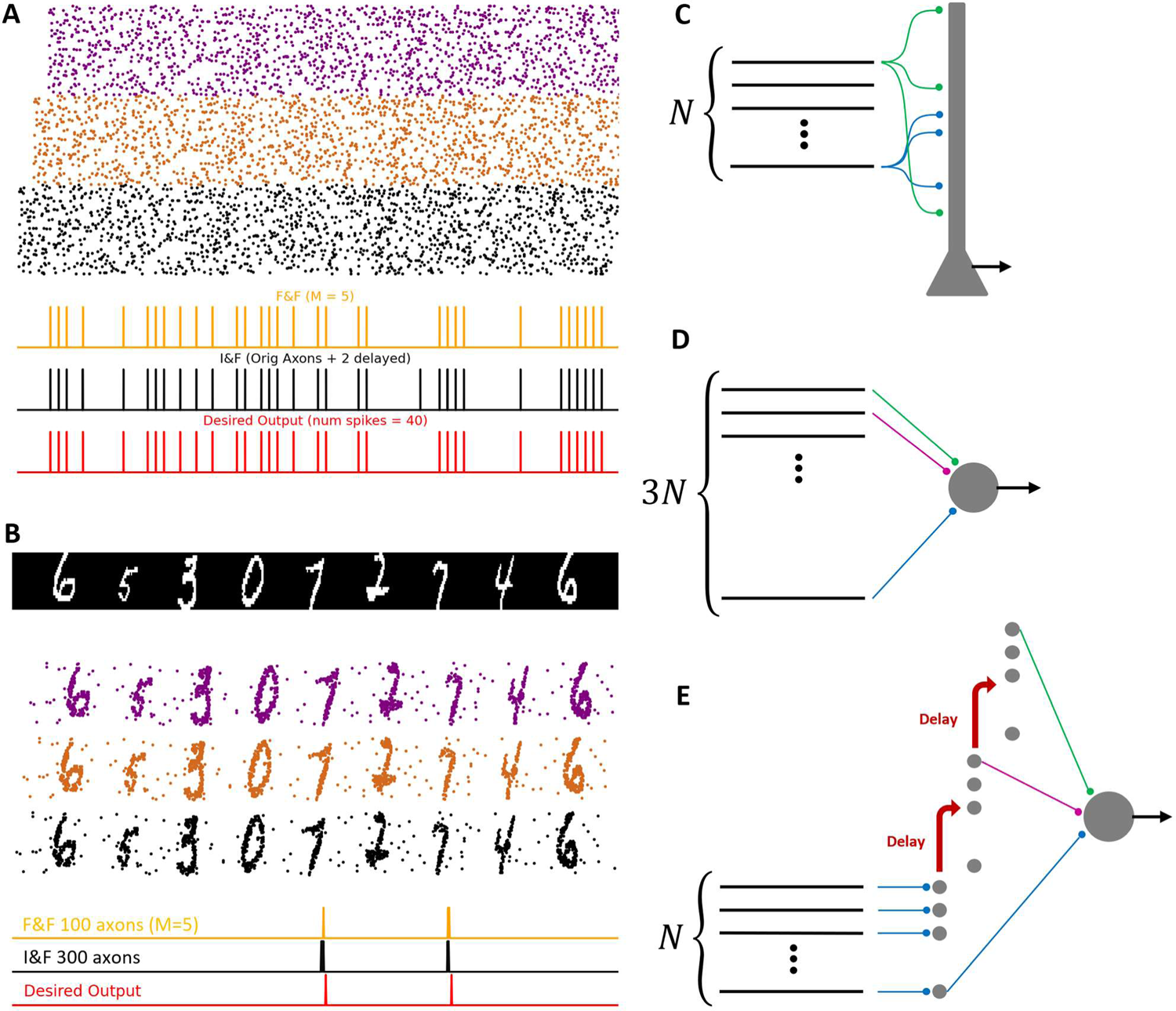
Multiple dendritic synaptic contacts per axon reduce the number of transmitting axons and/or decoding neurons. Illustration of three computationally equivalent implementations. (A) Input axonal raster composed of three sets of axons (black, brown and purple axons). Raster of the brown axons is a delayed version of the black raster, and the purple raster is delayed once more. In the first scenario, only the black axons are fed to a downstream F&F neuron, whose output is shown in orange (with 5 contacts per axon). In the second scenario, the full set of axons is fed into an I&F neuron (1 contact per axon), whose output is shown in black. The desired output spike train is shown in red. This panel demonstrates that an I&F neuron with 3N axons (with delayed input) replicates the precise timing capacity of the F&F model with N axons. (B) As in (A) but for the spatio-temporal MNIST digit recognition task. Here again an I&F neuron requires additionally delayed input axons to replicate the performance of the F&F neuron with N axons on the spatio-temporal MNIST digit recognition task. (C) Illustration of N axons that feed into a single F&F neuron model receiving multiple synaptic contacts. (D) For an I&F point neuron model to have identical memorization capacity as the F&F neuron illustrated in (C), the same information needs to be separated into 3N axons. (E) In order for an I&F point neuron to have identical spatio-temporal patterns separation capabilities with N axons as in (C), it needs to have a feed-forward “delay line” decoding network prior to being fed into the I&F “readout” neuron.

## Discussion

Cortical neurons typically connect to each other via multiple synaptic contacts. The computational implications of this apparent redundancy are unclear. To explore this question, we presented in the present study the Filter and Fire (F&F) neuron model that augments the commonly used Leaky Integrate and Fire (I&F) neuron model by incorporating in it multiple synaptic contacts per axon as well as the effect of dendritic filtering that transforms the incoming spike train to a set of multiple post synaptic potential (PSP) filters. Each filter corresponds to a particular synaptic contact, with varied time constants. This directly relates to the location-dependent filtering of the dendritic synaptic inputs by the dendritic cable properties^21,22^. In the F&F model, a single presynaptic spike impinging on several dendritic locations results in a somatic PSP composed of multiple temporal profiles. We termed this phenomenon “dendro-plexing” of the presynaptic spike (Figure 1).

We have demonstrated that the capacity of the F&F neuron to memorize precise input-output relationships is increased by a factor of ∼3 compared to that of the regular I&F. The capacity is measured as the ratio between the number of precisely timed output spikes and the number of incoming input axons. This ratio is ∼0.15 spikes per axon for the I&F case as was shown by Memmesheimer et al.^23^, and ∼0.45 spikes per axon for the F&F model (Fig. 2C). We showed in Figure 4, that the origin of this threefold increase in capacity is due to the fact that all possible PSP shapes are spanned by a subspace consistent of 3 basis shapes. This subspace sets the effective upper limit on the capacity when using any number of multiple synaptic contacts. i.e., even if using numerous synaptic contacts, they effectively serve as only 3 independent contacts.

Next (Fig. 3), we constructed a new spatio-temporal pattern discrimination task using the MNIST dataset and demonstrated that the F&F model can learn to detect single digits at well-above chance level performance on an unseen test set, whereas an I&F neuron model cannot learn the task at all, because in the specific way we chose to represent each digit – the task does not contain enough spatial-only information suitable for I&F neuron discrimination. Our specific task design was deliberately chosen to highlight this temporal aspect of pattern discrimination that is possible when taking into account the temporal filtering due to cable properties of dendrites. We show that multiple synaptic connections with different PSP profiles allow the neuron to effectively parametrize the temporal profile of the PSP influence of each pre-synaptic axon on the somatic membrane potential. This is enabled by modifying the weight of the various (multiple) contacts made between the axon and the post synaptic cell. We show that, for the range of PSP filters we have used, as well as for the range of somatic PSP in a detailed model of L5 cortical pyramidal cell (Fig. S2) all PSPs can be spanned by a 3 basis PSP filters, each with a different temporal profile. Taken together, this suggests that the F&F neuron model provides a low temporal frequency approximation to a spatio-temporal perceptron that assigns independent weights to each point in time. An alternative description that is mathematically equivalent is that the F&F model effectively bins the membrane integration time into 3 non-uniform time bins and can learn to assign independent weights for each temporal bin.

Our study demonstrates that even when considering highly simplified neuron model as used here, one that only implements the passive temporal filtering aspect of dendrites, a computational role of dendrites is unraveled for the seeming redundancy of multiple synaptic contacts between pairs of neurons in the brain. Dendrites therefore allow us to salvage some of the redundant connectivity and put those synaptic weight parameters to good use. This allows for increased memorization capacity, but perhaps more importantly to detect specific spatio-temporal patterns in the input axons. The spatio-temporal filtering properties of dendritic processing are prominently featured in our recent study describing how single neurons produce output spikes in response to highly complex spatio-temporal patterns^28^. It is important to note that there is an additional nonlinear amplification aspect in dendrites^29–32^ that we did not consider in this study and that is likely to provide additional computational benefits also in the context of multiple synaptic contacts, as we previously suggested^28^. We have therefore added in the present study a new perspective on the growing literature regarding the computational function of individual neuron ^29,30,33–44^

It is notable that it is typically believed that the phenomenon of multiple synaptic contacts is primarily attributed to noise reduction related to unreliable synaptic transmission, but our work strongly suggests that, from a computational standpoint, this is only a small part of the story. In fact, in principle each synapse can reduce its own unreliability by employing a multiple vesicle release (MVR) mechanism, as was also recently suggested by^12^. Here we suggest that unreliable synaptic transmission might play a computational role, a “feature” rather than being a “bug”. In Fig. 3**F** we show that synaptic “noise” might play a role that is similar to “dropout”^27^ or more precisely drop-connect^26^ mechanism which is commonly used in present day artificial neural networks paradigm. There, drop-connect is typically employed as a regularization technique that reduces overfitting and usually improves generalization.

The question of how trains of spikes represent information in the nervous system has been a long-standing question in neuroscience, since its inception. A major debate revolves around whether information is largely carried by firing rates averaged over relatively long time periods, or rather that precisely timed spikes carry crucial bits of information. Evidence for both alternatives has been found for both sensory systems and motor systems and, consequently, much theoretical work on this key topic has been conducted^23,45–62^. Since we have shown that dendrites and multiple synaptic connections per axon play a crucial role in decoding incoming spike trains and increase the neuronal repertoire in emitting precisely timed output spikes in response to spatio-temporal input patterns (and they might do this using a simple biologically plausible learning rule), we wish to suggest that dendritic hardware should be considered when discussing the question of the neural code and what information is transmitted via axons. Indeed, it was shown^63^ that a diversity of time constants helps increasing the computational repertoire of spiking networks.

Here we suggest that diversity of time constants, that helps in increasing the computational repertoire, already happens at the neuronal level. The fact that a single neuron can decode complex spatio-temporal patterns on its own and does not require a highly coordinated decoding network of neurons to extract temporal information from incoming spike trains, not only allows for potential “hardware” (wiring) savings as we illustrated in Figure 5, but also suggest that information might be ubiquitously encoded by precise spike times throughout the central nervous system. A single neuron can emit precisely timed output spikes in response to spatio-temporal inputs, as we showed here and was previously shown in simpler I&F models by ^23^. It is therefore not required to have a large and highly coordinated network of neurons to encode temporally precise patterns transmitted via axons. We believe that if already a single neuron can learn to generate such precise spike timing without relying on network mechanisms, this might increase the likelihood that neuronal information is encoded by precise spike timing as opposed to average firing rates (over relatively long periods of time) throughout the CNS.

Additionally, the three equivalent alternatives shown in Fig. 5**C,D****&E**, highlight the general scenario in which the evolutionary pressure to reduce axonal wiring in an ever-increasing brain volume^64,65^ could be solved by compressing information on a limited number of axons and by relying on sophisticated dendritic integration to decode these signals. Indeed, our study shows that employing multiple synaptic contacts between the axon and its postsynaptic neuron improves the decoding capability of spatio-temporal patterns by neurons and, at the same time, save “hardware” (reduced total axon length and/or number of neurons) at the cost of a small increase in single neuron complexity.

Lastly, in recent years, multiple groups around the world have started to generate dense reconstruction of rodent and human neuronal (cortical) circuits at the EM level, and report neuronal connectivity maps ^6,7,66–71^. We would like to suggest that analyzing these EM datasets, focusing on the number of multiple contacts and their locations on the dendritic tree, might shed additional light on the extent and role of multiple synaptic contacts between different cell types and brain regions, and hint to the possible “style” of information processing in these networks based solely on EM data. As illustrated in our work, if two neurons form a connection on distal dendrites, or if they form a connection on proximal dendrites, these will result in completely different time course of the somatic voltage and, therefore, on the temporal coding capabilities of the neuron. Indeed, simply reporting connectivity maps and even the size of post synaptic density (PSD) areas is not enough to determine the temporal influence of the connection on the post synaptic cell, as the dendritic location of the synapse is key, as was also shown by Rall ^21,22^ and recently also suggested in ^69^. Furthermore, due to the nonlinear amplification of dendrites, such EM-based data will be crucial in assessing whether two presynaptic neurons connect to nearby dendritic locations as, in this case, these synapses are much more likely to undergo nonlinear amplification (and might produce additional broad/slow NMDA or calcium-dependent temporal filters) if activated at similar times. This will add additional broader/long-lasting filters to neurons and increase their capability to learn to recognize spatio-temporal input patterns beyond what we have shown in this study.

## Methods

### F&F neuron simulation

A F&F neuron receives as input *N*_axons_ input axons, their spike trains will be represented by *X*_i_(*t*) = ∑_ti_ δ(*t* − *t*_i_) and they will be denoted by index *i*. Each axon connects to the dendrite via *M* contacts Each contact connects on the dendrite at a location that will be denoted by index *j*, and filters the incoming axon spike train with a specific synaptic kernel *K*_j_(*t*). The kernels are typical double exponential PSP shapes of the form *K*_j_(*t*) = *A* · (*e*^-t⁄rdecay,j^ − *e*^-t⁄rrise,j^) where A is a normalization constant such that each filter has a maximum value of 1, and τ_decay,j_, τ_rise,j_ are different for each contact, sampled randomly and independently for each contact from the ranges τ_decay,j_ ∈ [12*ms*, 30*ms*], τ_rise,j_ ∈ [1*ms*, 12*ms*]. Different kernel parameters represent a randomly connected axon-dendrite location.

The result of the kernel filtering of the corresponding input axons forms the contact voltage contribution trace *V*^i^ (*t*) = *X*_i_(*t*) ∗ *K*_j_(*t*) = ∑_t_ *K*(*t* − *t*_i_). There are overall a total of *M* · *N*_axons_ such contact voltage contributions traces. In vector notation we denote *V*_c_(*t*) = [*V*_c,1_(*t*), *V*_c,z_(*t*), ⋯, *V*_c,M·Naxons_(*t*)]. Each synaptic contact has a weight, *W*_j_. In vector notation we write *W* = [*W*_1_, *W*_z_, ⋯, *W*_M·Naxons_]. Each local synaptic response is multiplied by its corresponding weight to form the somatic voltage *V*_s_(*t*) = *W*^T^ · *V*_c_(*t*) = ∑_j_ *W*_j_ · *V*_c,j_(*t*). When threshold is reached, the voltage is reset, and a negative rectifying current is injected that decays to zero with a time constant of 15ms. Note that due to mathematical simplicity we do not impose any restrictions on synaptic contact weights, each weight can be both positive or negative regardless of which axon it comes from.

### I&F neuron simulation

The I&F simulation details are identical in all ways to the F&F neuron, except that all of its contact kernels are identical *K*_j_(*t*) = *K*_l∧F_(*t*) = *A*_l∧F_ · (*e*^-t⁄rdecay,l∧F^ − *e*^-t⁄rrise,l∧F^), where τ_rise,l∧F_ = 1*ms* and τ_decay,l∧F_ = 30*ms*.

### F&F learned weights visualization

In the F&F model a single input axon is filtered by multiple contact kernels *V*^i^ (*t*) = *X*_i_(*t*) ∗ *K*_j_(*t*). In order to display the effective linear model weights, we can group together all kernels that relate to the same axon *V*^i^(*t*) = ∑_ji_ *W*_ji_ · *X*_i_(*t*) ∗ *K*_ji_(*t*) = *X*_i_(*t*) ∗ ∑_ji_ *W*_ji_ · *K*_ji_(*t*). Therefore, the term ∑_ji_ *W*_ji_ · *K*_ji_(*t*) is a composite kernel that filters each axon. This is a function that can be visualized for each input axon, and therefore related directly to the input space.

### Capacity calculation details

To measure capacity, we sample *N*_axons_ random spike trains to serve as axons with a Poisson instantaneous firing rate of 4Hz for a period of 120 seconds. We randomly distribute *N*_spikes_ output spikes throughout the 120-second time period to generate *y* (*t*). For the sake of mathematical simplicity and not dealing with reset issues, we make sure that the minimal distance between two consecutive spikes is at least 120ms (4 · τ_decay,l∧F_). We bin time to 1ms time bins. We calculate *V*_c_(*t*) for the entire trace. Our task is to find a set of weights such that *y* (*t*) = φ(*W*^T^ · *V*_c_(*t*)), where φ(·) is a simple thresholding function. This is in essence a binary classification dataset with 120,000 timepoints (milliseconds), for each of those timepoints the required output of a binary classifier is either 1 (for time points that should emit an output spike), or 0 for all other timepoints. We have a total *M* · *N*_axons_ weights that need to fit the entire 120K sample dataset. We calculate the AUC on the entire 120K datapoint dataset. We declare that the fit was successful when AUC > 0.99. We repeat the procedure for various values of *N*_spikes_. The maximal value of *N*_spikes_ that we still manage to fit with AUC > 0.99 is termed *N*_spikes,max_ and the capacity measure is this number normalized by the number of axons used *ity*(*M*) = *N*_spikes,max_⁄*N*_axons_. Note that due to *N*_spikes_ ≪ 120,000, the capacity of the problem effectively doesn’t depend on the length of the time period we use, but by rather the number of spikes we wish to precisely time.

### Spatio-temporal MNIST task and evaluation details

The MNIST dataset contains 60,000 training images and 10,000 test images. The images are of size 28×28 pixels. We crop the images at the center to be 20×20 pixels, and binarize the values. We then convert the horizontal spatial image dimension (width) into a temporal dimension by uniformly warping the time such that 20 horizontal pixels will be mapped onto T milliseconds. T is the pattern presentation duration. The vertical spatial image dimension (height) is simply replicated 5 times so that 20 vertical pixels will be mapped onto 100 axons. The entire training set is the concatenated sequentially (in random ordering), with 70ms of zeros between every two patterns. We then sample spikes for every 1ms time bin according to each axon’s instantaneous firing rate to generate the raster for the entire training set. On top of that we add an additional background noise rate. The output ground truth is a single output spike 1ms after the pattern is presented for the positive class digit, and no spikes for negative patterns. The fitting of the model is identical to that described in the capacity sections. i.e., we wish to find a set of weights such that *y* (*t*) = φ(*W*^T^ · *V*_c_(*t*)), where φ(·) is a simple threshold function. In this case, we allow for wiggle room when evaluating performance on the test set. A successful true positive (hit) is achieved if at least 1 spike has occurred in the time window of 10ms around the ground-truth desired spike. A failed false positive (false alarm) is considered if a spike has occurred during the time window of 10ms around the end of the pattern presentation. We measure the classification accuracy under these criteria. We sometimes train only on a part of the training dataset and use only *N*_positivesamples_ as the positive class. In the regime of unreliable synaptic transmission with some synaptic release probability *p*, we sample the spikes of the input patterns once, and then present it *N*_epoch_ = 15 times, each time with random release probability samples for each presynaptic spike for each contact. In those cases, we perform the same procedure also for the test set (i.e., display the same pattern *N*_epochs_ = 15 times, each time with different synaptic release sampled for each presynaptic spike for each contact)

### Spatio-temporal Logistic Regression (LR) evaluation details

A spatio-temporal temporally sliding logistic regression (LR) model is a non-biologically plausible that is specifically used to serve as an aspirational model used for comparison for our proposed F&F neuron. It has a two-dimensional weight matrix *W*_LR_(*s*, *t*), where *s* spans the spatial dimention and *t* spans the temporal dimension. Mathematically, the equation describing the relationship between the input and the output of the model is given by *y*_LR_(*t*) = _φ(∑Naxons ∑_TLR _*W*_ (*s*, τ) · *X* (*s*, *t* − τ)). Note that this model has in total *N* weights (time is discretized into 1ms time bins here as well, as throughout this study).

## Data and materials availability

All code necessary to reproduce all results in this paper are available on GitHub via the link: https://github.com/SelfishGene/filter_and_fire_neuron All data and live scripts to reproduce all figures are available on Kaggle at the following link: https://www.kaggle.com/selfishgene/fiter-and-fire-paper

## Acknowledgments

We thank all lab members of the Segev and London Labs for many fruitful discussions and valuable feedback.

## Author contributions

D.B: Conceptualization, Methodology, Investigation, Visualization, Software, Validation, Data curation, Writing - Original Draft. S.S: Visualization, Software. I.S. and M.L: Conceptualization, Methodology, Writing - Review & Editing, Supervision, Resources, Funding acquisition.

## Supplementary Materials

**Fig. S1.**
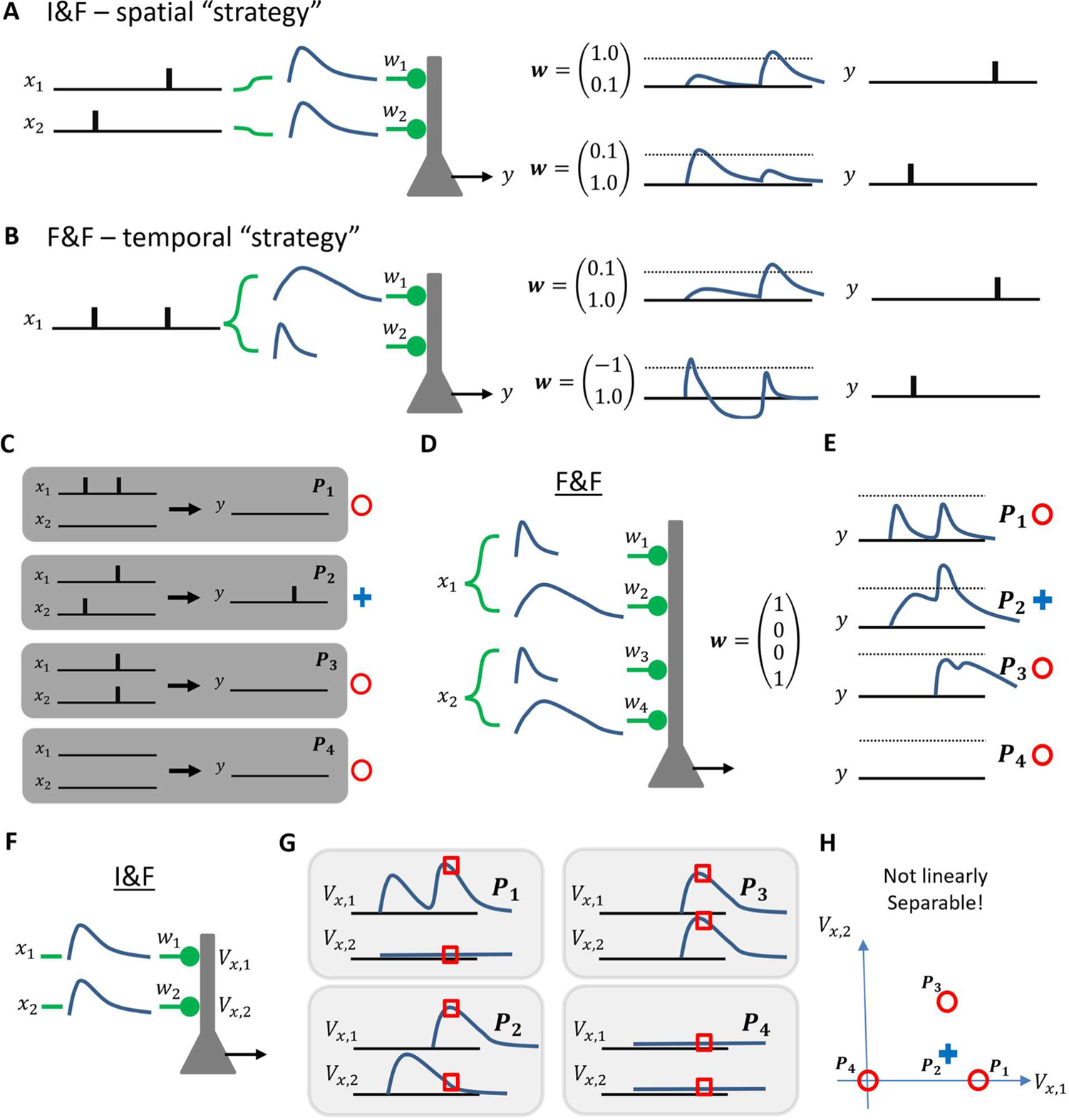
Toy illustration to highlight differences in computational capabilities between the F&F neuron and I&F neuron. (**A**) Illustration of the spatial strategy the I&F model employs. Left. Input axons x1, x2 are fed into an I&F model to emit an output y. Right, two output for different weight configurations that result in two different output patterns y. Dashed line represent spike threshold. (**B**) Illustration of the temporal strategy the F&F model employs. Left. A single input axon x1 is fed into an F&F neuron with two different PSP filters to produce output y. Right, two output for different weight configurations that result in two different output patterns y. note that although the output patterns are identical to those in (A), this was achieved by receiving a single input axon. (**C**) Example of a temporal task we wish to teach the two neuron models. For the four different input patterns (x1, x2) we require to produce the output y on the right. (Patterns are denoted as P1, P2, P3, P4). Only P2 is required to emit an output spike (marked with blue plus sign), the other patterns are required to emit no spikes (marked with red circle) (**D**) Illustration of a F&F model with fast and slow PSP filters for each input axon that can solve the task. (**E**) The somatic responses for each pattern with the weights vector w that solves the problem. The dotted lines represent the spike threshold. (**F**) Illustration of the I&F neuron model, its inputs and synaptic contact voltage contribution illustrated. (**G**) The normalized unweighted synaptic contact contribution voltage traces for all 4 patterns. The red squares mark the point in time where an output spike is expected. (**H**) The I&F neuron model cannot classify the output patterns as the four patterns are not linearly separable in the dendritic representation space (Vx,1 Vx,2) as seen in the illustration, and therefore a satisfactory weight vector for the synapses does not exist.

**Fig. S2.**
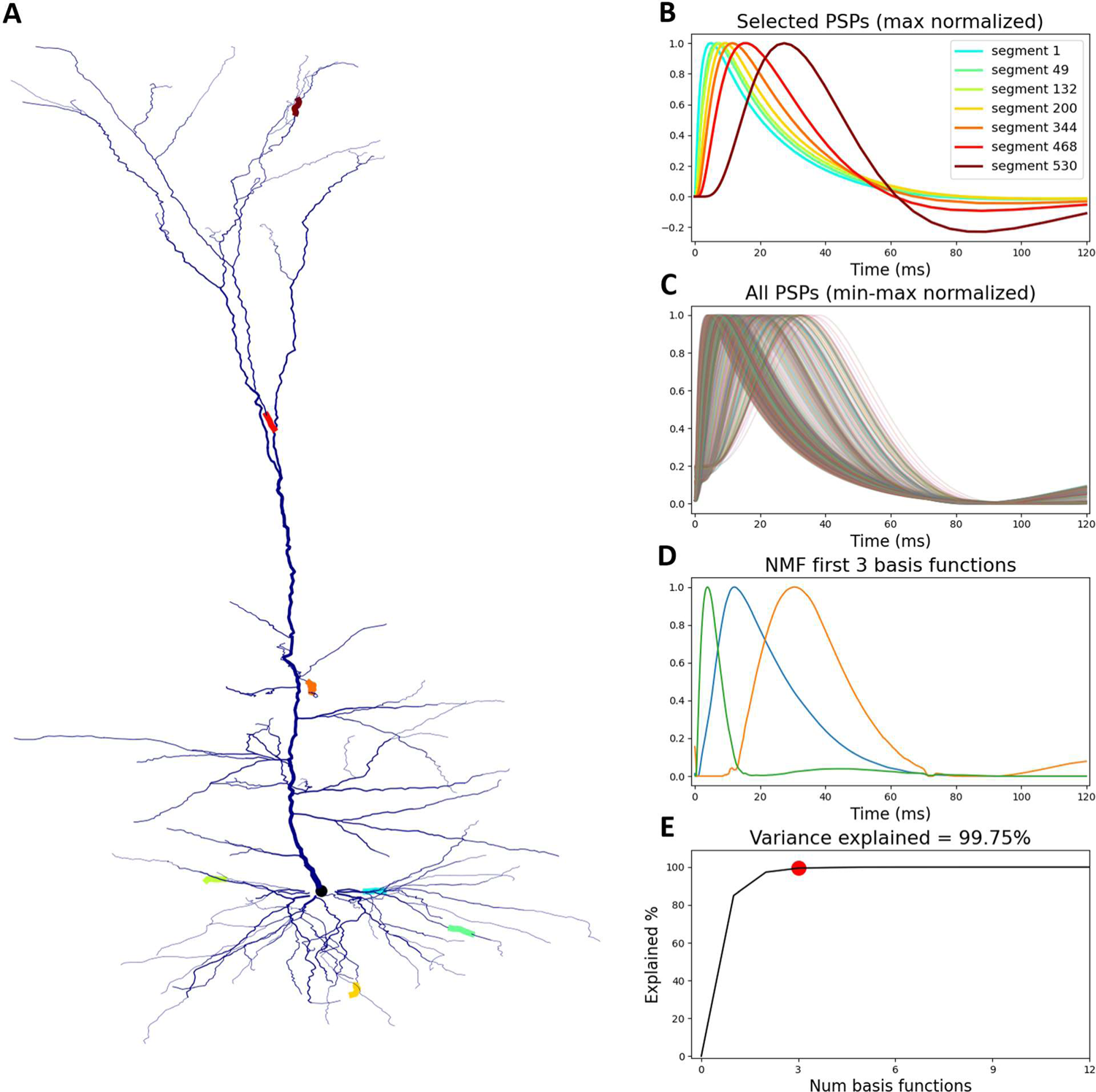
The dendritic filters of a reconstructed L5PC detailed biophysical model are also spanned by a three-dimensional basis set of PSPs as does our simplified F&F neuron throughout the paper (A) The morphology of a L5PC that was biophysically modeled in detailed ^72^ with highlighted dendritic segments. (B) Normalized somatic PSPs in response to excitatory synaptic input at the highlighted dendritic segments in (A). Note that in some of the EPSPs there exists a small hyperpolarization that is due to nonlinear potassium channels at the soma. (C) Normalized somatic EPSPs from all dendritic segments in the modeled L5PC. The traces are normalized such that minimal value in the time window is 0 and maximal value is 1. (D) First three basis functions of the non-negative matrix decomposition of all EPSPs shown in (C). NMF instead of SVD was used for simplicity of interpretation and visualization purposes. These filters look very similar to those shown in Fig. 4D. (E) Cumulative variance explained by each basis component demonstrating that the three basis functions shown in (D) can span all EPSPs shown in (C).

